# Live-trapping of rodents in urban green spaces across Los Angeles

**DOI:** 10.1101/2023.03.13.531394

**Authors:** Anthony R. Friscia, Sarah Helman, Molly Maloney, Alexandra K. Molina-Echavarria, Sarah Nugen, Nihal Punjabi, Isobel Tweedt, Jessica W. Lynch

## Abstract

Urban green spaces have the potential to function as multi-benefit spaces, for both human enjoyment and sustaining native wildlife populations. In our study, we trapped for small nocturnal mammals across a series of Los Angeles green spaces. Our results show that native rodents are only present in habitats that support native vegetation; in particular we highlight the native rodent biodiversity on Sage Hill, a coastal sage scrub remnant on the UCLA campus. Other urban parks that are composed of non-native grasses and non-native shrubbery yielded only invasive species of rodents, including Brown Rats (*Rattus norvegicus*) and House Mice (*Mus musculus*). Our study points to the ability of renovated green space in Los Angeles to support native fauna. In addition, our study demonstrates some of the difficulties in doing trapping studies in heavily urbanized environments.

---

In our study we used live trapping to investigate whether green spaces across Los Angeles support native rodent populations, and what factors might affect the presence of native versus introduced small mammal species. “Green spaces” are often thought of, and sold to the public, as tools for conservation of local fauna in metropolitan areas (Ofori et al. 2018). The 2016 Los Angeles Countywide Comprehensive Parks and Recreation Needs Assessment, written by Los Angeles County Department of Parks & Recreation^1^, states that Los Angeles parks built or renovated in the future should be ‘multi-benefit’, not only for human enjoyment but also contributing to the enhancement of regional sustainability and provision of ecosystem services. Testing whether local fauna use metropolitan green spaces is key to future planning, both as a tool for conservation and for popularizing conservation efforts.

Sage Hill at the University of California, Los Angeles (UCLA) is the last patch of natural coastal sage scrub vegetation on campus^2^, and in fact in the entire area of west Los Angeles south of Sunset Boulevard. It has been promoted as a haven not just for the natural flora of the area, but also the fauna. In 1996 a UCLA Geography course^3^, performed an inventory of the vegetation, birds, herpetofauna, arthropods, and mammals at the site. In their small mammal inventory, both native and invasive rodents were trapped at Sage Hill, including two Big-Eared Woodrats (*Neotoma macrotis*), one Brown Rat (*Rattus norvegicus*), and two House Mice (*Mus musculus*), as a result of trapping efforts of 32 traps on two different nights. The 1996 trapping success rate varied from 6.25% to 15.6% trap success on the two nights of capture.

In comparison, similar trapping methods and efforts at a continuous area of similar habitat, Leo Carrillo State Beach, yield a lower number of species but much higher trapping success, and a higher proportion of native species trapped^4^. Another local study that catalogued small mammals was carried out in 2000 in the Baldwin Hills area of Los Angeles by Dines^5^. This area retains native vegetation and also contains large preserved green spaces, e.g., Kenneth Hahn State Park, as well as much of the current oil exploitation in the Los Angeles Basin (see Fig. 1). The Baldwin Hills study included live-trapping of nocturnal small mammals. Four nights of trap-lines were set, and four species of rodent were trapped, including one invasive species, the house mouse, and three native species to southern California, the California Pocket-Mouse (*Chaetodipus californicus*), the California Vole (*Microtus californicus*), and the Big-Eared Woodrat, with a success rate for trapping efforts at about 20%. The current project began as an effort to provide a new inventory of the small mammals at Sage Hill twenty years after the 1996 survey, and later expanded to inventory green spaces across urban Los Angeles for nocturnal small mammals.

**Fig. 1.**
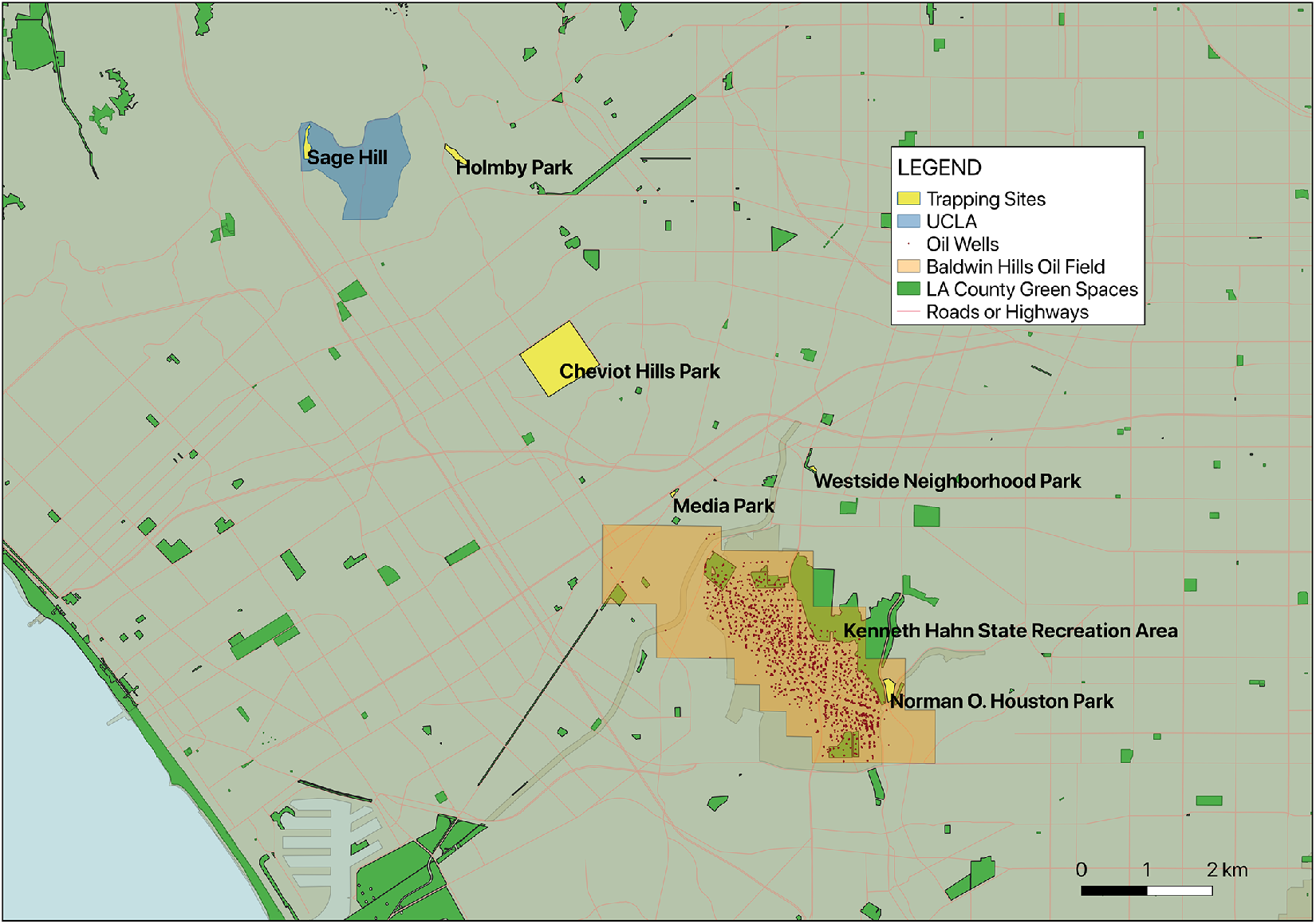
GIS Image of western Los Angeles. Location of trapping sites and other localities mentioned in the text are shown. North is towards the top. Image created using QGIS (QGIS Development Team, 2020) and open-source L.A. city and county GIS datasets.

## MATERIALS AND METHODS

In a pilot study to catalogue small nocturnal rodent species presence on Sage Hill, on the northwestern corner of UCLA campus, a trapping program was carried out in Spring 2015 and in Spring 2016 by the first author (ARF) and students for the UCLA GE70C seminar, “Evolution: A Naturalist’s Perspective” (a full list of students from the 2016 course is provided in Acknowledgments). Sage Hill is bounded on the west by Veteran Avenue and on the east by the residence halls along De Neve Drive. Three nights of trapping were performed on Sage Hill during each class. In each case, fifty traps were set in 3 trap lines, mainly on the south end of Sage Hill, starting from the bottom of the hill and ending at the top of the hill near the fencing that separates Sage Hill from the residence halls. Traps were set about 3 m apart, although this was modified based on local terrain and vegetation. 7.62×8.89×22.86cm Sherman-style live traps were used to allow for capture-and-release of animals (Hoffman et al. 2010). Trap size was selected to allow for the largest rodents (rats) to fit but to prevent larger animals (e.g., rabbits or foxes) from being able to enter. Traps were baited with a combination of oats (Hoffman et al. 2010; Brehme et al. 2011), peanut butter (Hoffman et al.) and diced apple (Frye et al. 2015). Raw cotton was added to each trap to allow for nesting and help alleviate any effects of cold weather at night (Wilson et al. 2007). All traps were closed during the day, baited just before sunset, and checked soon after sunrise. All specimens were handled according to UCLA Office of Animal Research Oversight protocols (protocol #2015–052–01). Upon verification of an occupied trap, the trap was inverted into a clear, plastic bag, which allowed for identification to species (Reid 2006) prior to release. While marking or tagging of trapped individuals would have been ideal, this was not possible in this pilot study.

The success in the pilot study of documenting native small mammals on UCLA campus became the impetus for a larger study to investigate the presence of rodents, especially native species, at other “green spaces” across the Los Angeles urban landscape which include little, if any, native vegetation. In 2018 trapping was carried out at Los Angeles City Parks (with permission from park officials) and trapping also was repeated at Sage Hill. Trapping was performed under scientific collecting permit SC-13720 from the California Department of Fish & Wildlife and all animal handling carried out by ARF in accordance with procedures laid out by the UCLA Research Safety & Animal Welfare Administration (ARC# 2015-052-01). Funding for the 2018 study was through UCLA Grand Challenges: Sustainable Los Angeles, under the Urban Ecology of Los Angeles Mammals grant. Several undergraduate students at UCLA took part in the trapping effort as student interns and volunteers.

Trapping occurred at five Los Angeles city parks as well as Sage Hill (Fig. 1 and Table 1). Parks were chosen based on the presence of enough vegetation to allow for placement of traps. Parks ranged in size from 0.68 ha to 10.9 ha, and other than Sage Hill, were mainly groomed spaces with lawns and introduced shrubbery and very little native vegetation or ground cover over cut-grass height, as is typical of Los Angeles parks. We performed two trapping sequences of three nights each at each location. One set of trapping sequences spanned late spring through early summer of 2018, and the other the fall of 2018. Each night, 30 traps were set in trap-lines, with traps set in the same places each night. Trapping protocol in 2018 was the same as described above for the 2015-16 Sage Hill trapping effort. More information about specifics of where traps were set is available upon request.

**Table 1.**
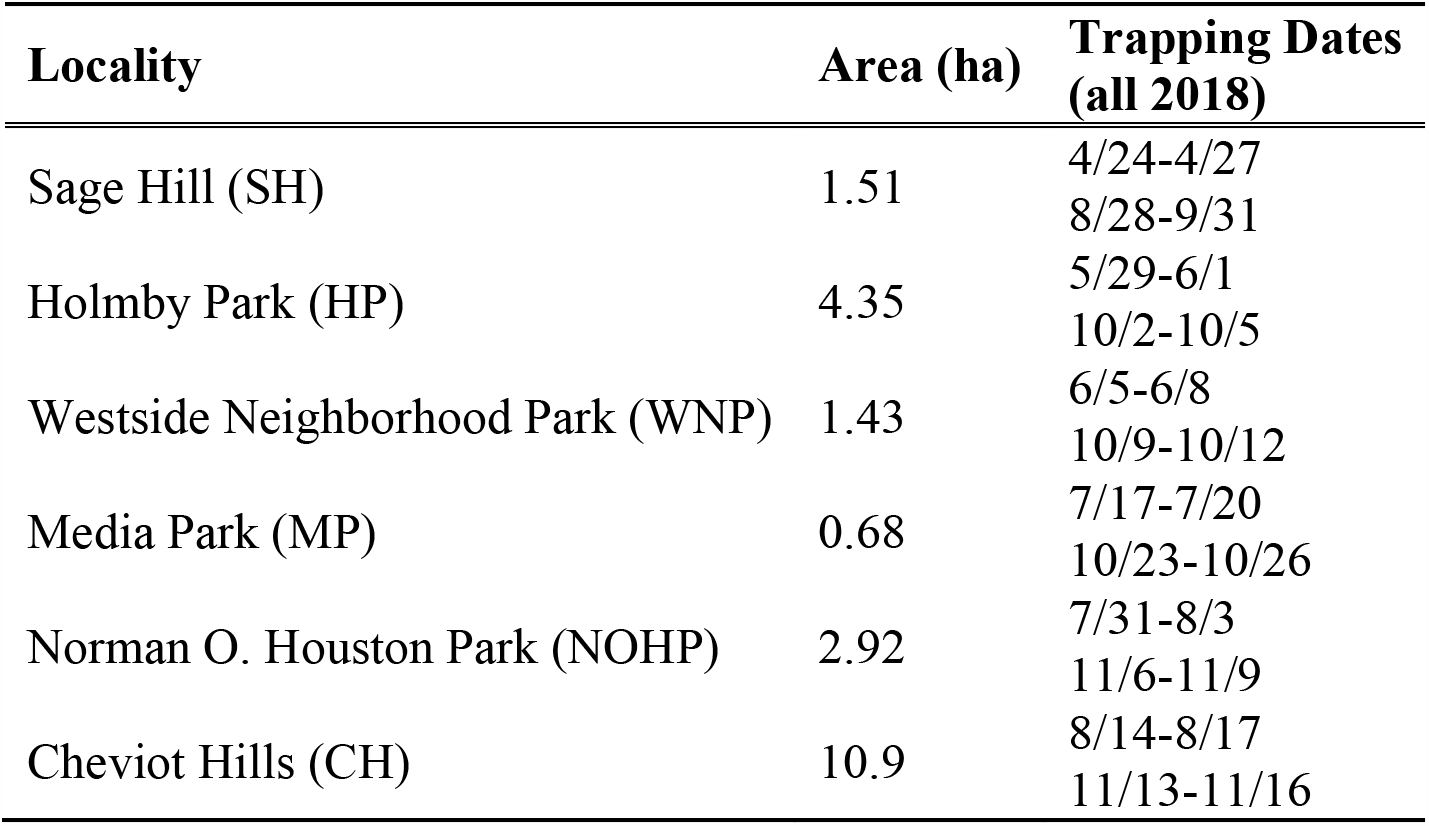
Trapping locations, areas and dates in 2018.

## RESULTS

In Fall 2015, three nights of trapping at Sage Hill did not result in the capture of any small mammals (see discussion). In 2016, over the course of 150 trap nights (50 traps over 3 nights) at Sage Hill, nine animals were captured (a success rate of 6%). Five different species were captured, out of which four were southern California native species: Deer Mouse (*Peromyscus maniculatus*), trapped on 2 nights; California Vole, trapped on two nights; Big-Eared Woodrat, captured on three nights (but may have been the same individual on different nights, given the woodrat nest near to the trapping site); and California Pocket-Mouse (*Chaetodipus californicus*), trapped on one night. The diurnal California Ground Squirrel (*Otospermophilus beecheyi*) commonly are observed on Sage Hill, bringing the total number of native rodents up to five. The only invasive species captured was the house mouse. Neither the black rat nor the brown rat was captured, although rats are commonly observed in other parts of the campus.

Across all Los Angeles green space sites and all trapping sequences in 2018, there were a total of 1080 ‘trap nights’, and a total of 29 captures, for a trapping success rate of 2.7% (see Table 2). The low number of ‘free lunches’ during the study, where bait was missing without the trap being sprung, indicate that the traps were typically sprung if an animal entered them. Trapping success rate varied across sites and trapping sequences. For example, trap success rate at Cheviot Hills was 11% in the spring, but 0% in the fall. Although lunar cycle is known to affect small mammal activity patterns (Owen et al., 2021), we did not attempt to correct for this as most parks were lit throughout the night and there seemed to be no correlation of trapping success and lunar phase. Of the animals captured throughout the main study across Los Angeles parks, only two were native California species (Big-Eared Woodrat and Deer Mouse), and these were both captured at Sage Hill (see Table 2). No native species were captured in city parks, not even at Holmby Park, a park of a similar distance to the Santa Monica Mountains as Sage Hill, nor at Norman O. Houston Park, a park very close to Baldwin Hills, an area with extensive native vegetation where a previous trapping study revealed several species of native small mammals.

**Table 2.** Trapping success and animals trapped by locality in main study.

| **<u>Locality</u>** | **<u>Trapping</u>**<br>**<u>Dates</u>** | **<u># of</u>**<br>**<u>Animals</u>**<br>**<u>captured</u>** | **<u>Black</u>**<br>**<u>Rat</u>** | **<u>House</u>**<br>**<u>Mouse</u>** | **<u>Big-</u>**<br>**<u>Eared</u>**<br>**<u>Woodrat</u>** | **Undetermined** |
| --- | --- | --- | --- | --- | --- | --- |
| SH | 4/24-4/27 | 1 |  |  | 1 |  |
| HP | 5/29-6/1 | 3 |  | 3 |  |  |
| WNP | 6/5-6/8 | 0 |  |  |  |  |
| MP | 7/17-7/20 | 2 | 2 |  |  |  |
| NOHP | 7/31-8/3 | 2 |  | 2 |  |  |
| CH | 8/14-8/17 | 10 | 9 | 1 |  |  |
| SH | 8/28-8/31 | 1 |  |  |  | 1 |
| HP | 10/2-10/5 | 1 | 1 |  |  |  |
| WNP | 10/9-10/12 | 0 |  |  |  |  |
| MP | 10/23-10/26 | 0 |  |  |  |  |
| NOHP | 11/6-11/9 | 9 |  | 9 |  |  |
| CH | 11/13-11/16 | 0 |  |  |  |  |
| **TOTAL** |  | **29** | **12** | **15** | **1** | **1** |

Abbreviations for localities follow those in Table 1.

The species captured at Sage Hill across the different years of study were the more common nocturnal small mammals in Southern California. The Deer Mouse is probably the most widespread native mouse in North America (Reid 2006) (but see Kalkvik et al. 2012 for alternate taxonomy). The cover provided by the vegetation at Sage Hill, and the presence of some shrubbery and trees provides the perfect habitat for the mice. It is unknown whether the mice range across the entire campus.

The Big-Eared Woodrat was captured every night in the 2016 trapping, and was captured again in 2018. A woodrat nest is present near the area where it was consistently captured, and its presence has been noted at Sage Hill before. Our trapping area at Sage Hill probably only has enough acreage to support one woodrat individual (Innes et al. 2012), although the entire acreage of Sage Hill has a number of woodrat nests.

The California Vole is another common species, present throughout wetter habitats in California (Kalkvik et al. 2012). The grass cover in the trapping area at Sage Hill makes it a suitable habitat. It would be interesting to determine if voles are prey species for local predators as they are a key prey species in native habitats (Kalkvik et al., 2012).

One small native mouse caught at Sage Hill was unable to be identified before it escaped handling. A tentative identification is a California Pocket-Mouse. The California Pocket-Mouse has been recorded in the Santa Monica Mountains, so it would not be surprising to find it at Sage Hill, but its presence should be confirmed with further trapping.

Invasive species captured at Sage Hill across our study were limited to one House Mouse in 2016. Across all the Los Angeles green spaces with non-native vegetation that we sampled for trapping small mammals, we captured only invasive species, including rats and house mice (Table 2).

## DISCUSSION

Our 2016 trapping effort done as part of a UCLA undergraduate class project demonstrated the presence of multiple native small mammal species on Sage Hill, a relatively isolated patch of native habitat. Prior to trapping, we expected to capture a number of individuals of invasive species, like the house mouse and the black or brown rat which are commonly seen across campus, and are often pests in buildings. The fact that we did not catch many invasive species at Sage Hill speaks to a possibly important role for the patches of natural habitat in urban settings. They could act as important refuges for native species. Further trapping should be done elsewhere on campus to find out if the native rodent species are found across UCLA campus, or only in this preserved habitat. This could be done with the help of campus pest control, who could provide information on the individuals they capture. This also raises the question of how much connection Sage Hill has to larger populations of native species in the Santa Monica Mountains. The only way to address this would be a more in-depth study involving the population genetics of Sage Hill and nearby mountain areas and/or tracking studies of individuals found at Sage Hill.

Studies conducted in cities around the world have found that native species compose only a small minority of the flora found within urban green spaces (Bolger et al. 1997; Figueroa et al. 2018). This makes Sage Hill, where California natives make up nearly 100% of the flora in some vegetative patches, especially unique^6^. In general, species richness is reduced in areas of extreme urbanization (McKinney 2008). Invasive species, which spread more easily in urban settings, can cause significant ecological imbalances, disrupting beneficial ecosystem services, such as energy, nutrient, and water cycling, that humans often depend upon (Charles & Dukes, 2007). Near-natural habitats are better equipped to resist the establishment of invasive species (Gong et al. 2013). Therefore, protecting and expanding relatively undisturbed habitats, like Sage Hill, is crucial for improving the biodiversity and ecosystem services available in urban settings. Our study suggests that, despite being far removed from other coastal sage scrub habitats, this relatively small area was still better suited to harbor native rodents than larger, more artificial green spaces. In addition, Sage Hill’s location on the UCLA campus makes it an invaluable resource for the academic community. The site resembles much of the surrounding area prior to urban development^7^, and thus allows for an understanding of southern California’s ecological history. Sage Hill already has been the site of multiple undergraduate class projects, and the present study afforded several undergraduate students early research experience. Protecting Sage Hill not only makes invaluable knowledge accessible to students and researchers, but also preserves a model to inform the design of future Los Angeles parks.

Our Fall 2015 trapping effort at Sage Hill yielded no captures of small mammals, but the same amount of trapping effort in Spring 2016 yielded four species of native taxa. The main difference between the two times was that in fall, all the non-native grasses had been cut, so there was very little cover in the area. Cover, native or otherwise, is an important driver of rodent occupancy of an area.

We expected some relationship between the presence of native animals to the size of the park or proximity to native habitats (Bolger et al., 1997), however neither relationship was observed. Holmby Park is situated a similar distance to the Santa Monica Mountains as Sage Hill (∼1 km), but no native species were captured. Similarly, Norman O Houston Park is in the Baldwin Hills, much of which are a relatively large green space with native flora and native rodent species, and while the Park had the overall highest trap rate success of the study (6.1%), only invasive species were captured. This park is across a major thoroughfare (La Brea Avenue) from the main area of Baldwin Hills, suggesting that roads with high traffic volume may be enough to block movement of native rodents here (Riley et al., 2006).

In Los Angeles many city parks have no vegetation; a common type of park is just playground equipment on artificial surfaces. The parks that were ‘urban green spaces’ chosen for this study had some kind of vegetation, either native or, more commonly, introduced. Some parks did have small, decorative patches of native plants (e.g., Westside Neighborhood Park), but they weren’t contiguous and the city parks were all dominated by groomed lawns or bare ground and non-native shrubbery. In our study, native rodent species were never captured in these green spaces without native vegetation.

The overall low trapping success rate at the city parks could be attributed to a number of factors. Food in the form of trash was readily available. In addition, most city parks had baited kill traps set by park management for pest control. As described in Desvars-Larrive et al. (2018), trapping difficulty for urban rats can be heightened by food abundance and predator presence (including domestic pets), among other variables such as rodenticide use. Small rodent abundance is also known to fluctuate annually (Blaustein, 1981), and in Southern California there tend to be low population sizes in winter and higher population sizes in spring and summer during the breeding season, with summer to early fall the maximum density^8^. In urban environments partnerships with pest control agencies might help yield data unavailable through traditional conservation-based trapping; this avenue is being explored by other members of our research group (Kelty, in prep).

Trapping in urban areas leads to a number of other difficulties as well. Although we had permission to trap from the head ranger of the LA City Parks, that information did not always make it to down to ‘front line’ workers. On one of our first trapping nights, at Holmby Park, we left our traps out (but closed) during the day and when we came to bait them in the evening they were all gone. A groundskeeper had taken all of them (and destroyed a few) thinking that they were illegally placed. They were returned to us only after explaining to him what we were doing, and making a call to the head ranger. After this we always removed traps during the day, but replaced them in the same locations each night. Park patrons were also wary of our work. They worried that we were using poisoned bait that could be obtained by children or pets. A key part of the work of rodent trapping surveys is the public education component (Desvars-Larrive et al.) and our team engaged in many conversations with the public about our project.

## CONCLUSIONS

“Green spaces” are often portrayed as wildlife refuges in urban environments. This can only be verified through censusing of wildlife in these spaces. Although birds and some insects would easily move between green spaces and between urban and natural environments, small mammals would almost certainly be more constrained in their movements. In our study, invasive rodent species dominated non-native green spaces across the city. Although preliminary, this study also shows that native small mammal species will take advantage of preserved habitats within an urban environment. In addition, they might outcompete invasive species in these preserved patches, highlighting the importance of native vegetation in preserving local small mammal biodiversity, a key observation that should be considered in future park creation and renovation. We especially want to highlight the unique characteristics of Sage Hill on UCLA campus, and the importance of protecting this native area. We also want to emphasize the difficulties in trapping in a highly urbanized landscape, as well as the utility of undergraduate coursework that involves local students in cataloguing biodiversity on an urban campus and in the surrounding urban landscape. Future studies will work to increase the sample size as well as utilize proper mark-and-recapture and population genetic methods to better characterize the rodent population in LA green spaces.

## ACKNOWLEDGEMENTS

We thank the other members of the LA Mammals research group, including Chris Kelty, Jamie Lloyd-Smith and Katie Prager for their support. A.R.F. thanks the students in the 2015 GE70 spring seminar, “Evolution: A Naturalist’s Perspective”: Ananya Bhargava, Jocelyn Correa, Rudy Diaz, Elisa Escamilla, Lauren Finkle, Nico Flores, Jeanette Gomez, Benjamin Kesner, Morgan Kutzner, Eugene Lee, Jehu Lee, Gaby, Meza, Viktor Nunes Peinemann, Tofunmi Oyebolu, Yesenia Perez, Jhem Quintana, Ryan Teng, Owen Weitzel, Tiffany Yang, Kerri Yund. We thank the City of Los Angeles, Department of Recreation and Parks for trapping permission. Jim Dines and Kathy Molina provided discussion and logistical support.

## Footnotes

1 Los Angeles County Department of Parks & Recreation. 2016. Los Angeles Countywide Comprehensive Parks and Recreation Needs Assessment. Placeworks, Available on-line: https://lacountyparkneeds.org/wp-content/uploads/2016/06/FinalReport.pdf

2 Mattoni R., Angulo A., Burnam J., Chalekian J., Chen J., Cortez N., Duvernay E., Farris C., Fry D., Hill R., Hillway K., Post A., Pun, K., Scholnick J., Sway D., Varvel, A., Wilson G., and Yancey C. 1997. Biological Assessment: Coastal Sage Scrub at University of California, Los Angeles. Geography 123: Bioresource Management UCLA Department of Geography, Winter 1996, Longcore T., Mattoni R., Maldonado J., Hertel F., Scow J., editors. Los Angeles, CA.

3 *Ibid*

4 *Ibid*

5 Dines J.P. 2001. Mammals of the Baldwin Hills. The Biota of the Baldwin Hills: An Ecological Assessment. Pp. 127–141 in Community Conservation International and Natural History Museum of Los Angeles County (Molina KC, ed.), Los Angeles, USA.

6 Mattoni R., Angulo A., Burnam J., Chalekian J., Chen J., Cortez N., Duvernay E., Farris C., Fry D., Hill R., Hillway K., Post A., Pun, K., Scholnick J., Sway D., Varvel, A., Wilson G., and Yancey C. 1997. Biological Assessment: Coastal Sage Scrub at University of California, Los Angeles. Geography 123: Bioresource Management UCLA Department of Geography, Winter 1996, Longcore T., Mattoni R., Maldonado J., Hertel F., Scow J., editors. Los Angeles, CA.

7 *Ibid*

8 Impact Sciences, Inc. 2005. Draft Assessment and Survey of Mammals within the Newhall Ranch Specific Plan Area. Los Angeles County, California, USA.

